# Treatment-seeking for vaginal fistula in sub-Saharan Africa

**DOI:** 10.1101/623520

**Authors:** Samson Gebremedhin, Anteneh Asefa

## Abstract

**Background:** There is dearth of data regarding the treatment-seeking practice of women living with vaginal fistula. The paper describes the health-seeking behaviour of fistula cases in the sub-Saharan Africa (SSA) where the burden of the problem is high.

**Methods:** We analysed the data of 1,317 women who ever experienced vaginal fistula, extracted from 16 national Demographic and Health Surveys carried out in SSA between 2010 and 2017. The association between treatment-seeking and basic socio-demographic characteristics assessed via mixed-effects logistic regression and the outputs are provided using adjusted odds ratio (AOR) with 95% confidence intervals (CI).

**Results:** Two-thirds (67.6%) of the women encountered the fistula soon after delivery implying obstetric fistula. Fewer identified sexual assault (3.8%) and pelvic surgery (2.7%) as the cause. In 25.8% of the cases clear-cut causes couldn’t be ascertained and excluding these ambiguous causes, 91.2% of the women had obstetric fistula. Among those who ever had fistula, 60.3% (95% CI: 56.9-63.6%) sought treatment and 28.5% (95% CI: 25.3-31.6%) underwent fistula-repair surgery. The leading reasons for not seeking treatment were: unaware that it can be repaired (21.4%), don’t know where to get the treatment (17.4%), economic constraints (11.9%), healed by itself (11.9%) and embarrassment (7.9%). The regression analysis indicated, teenagers as compared to adults 35 years or older [AOR=0.31 (95 % CI: 0.20-47)]; and women devoid of formal education when compared to women with any formal education [AOR=0.69 (95% CI: 0.51-0.93)], had reduced odds of treatment-seeking. In 25.9% of the women who underweight fistula-repair surgery, complete continence after surgery was not achieved.

**Conclusion:** Treatment-seeking for fistula remains low and it should be augmented via mix of strategies for abridging health-system, psycho-social, economic and awareness barriers.

## Background

Vaginal fistula is an abnormal communication between vagina and adjacent tubular structures – usually bladder and rectum – leading to continuous leakage of urine or faeces through vagina [1]. While virginal fistula has multiple causes, the leading aetiology leading to more than 90% of the global burden of fistula is injury associated with prolonged obstructed labour. This is commonly referred to as obstetric fistula [2]. Other less frequent causes include sexual assault, iatrogenic surgical damage, malignancy and radiation [1].

Vaginal fistula is a debilitating condition that has pervasive consequences on women’s psychological, social, physical and economic wellbeing. Without surgical intervention women with fistula face lifetime embarrassment, isolation, social stigmatization and marital separation [3-5]. Furthermore, they are prone to develop chronic vaginal and urinary tract infections, renal failure, pelvic inflammatory disease and amenorrhea [1,6].

The World Health Organization (WHO) estimated that annually 50,000 to 100,000 women worldwide develop obstetric fistula. While the problem had already been eliminated in the western world, up to three million women in sub-Saharan Africa (SSA) and Asia, are suffering from obstetric fistula [7,8]. The lifetime prevalence of vaginal fistula in SSA is as high as 3 cases/1,000 women of reproductive age and the figure exceeds 5 cases/1,000 women in many countries including Uganda, Kenya, Ethiopia and Tanzania [2].

The most effective way to treat virginal fistula is surgical closure of the defect [1]. However, in many high-burden countries, access to skilled professionals capable of repairing fistula is limited and very few hospitals provide the service. Further, as most of fistula cases are from impoverished and marginalized segments of population, treatment-seeking is likely to be hindered by psychosocial, economic and other contextual factors [9].

Multiple facility-based qualitative studies and crude estimates suggested that timely treatment-seeking for fistula is generally low and globally as high as two million cases remain untreated [6,10-12]. Yet, empirical quantitative evidence is limited possibly due to the fact that vaginal fistula is a rare event in statistical sense and recruiting adequate cases through community-based surveys is usually infeasible. Recently, Demographic and Health Surveys (DHS) surveys collect treatment-seeking related information in many developing countries but the findings have not been reported possibly due inadequacy of sample size. This study pools cases from multiple and recent DHS surveys conducted in the SSA region and describes treatment-seeking practices at regional level. Secondary indicators including reasons for not seeking care and success rate for fistula-repair surgery are also presented.

## Methods

### Study design and eligibility criteria

The study is conducted based on the secondary data of 16 DHS conducted in the SSA region between 2010 and 2017. Nineteen surveys that gathered fistula-related information were considered eligible. However, three surveys (Togo 2013-14, Senegal 2017, Burundi 2016-17) did not collect treatment-seeking related data and hence excluded from analysis. The final dataset comprised the data of 1,371 women with fistula from 14 countries (Figure 1).

**Figure 1:**
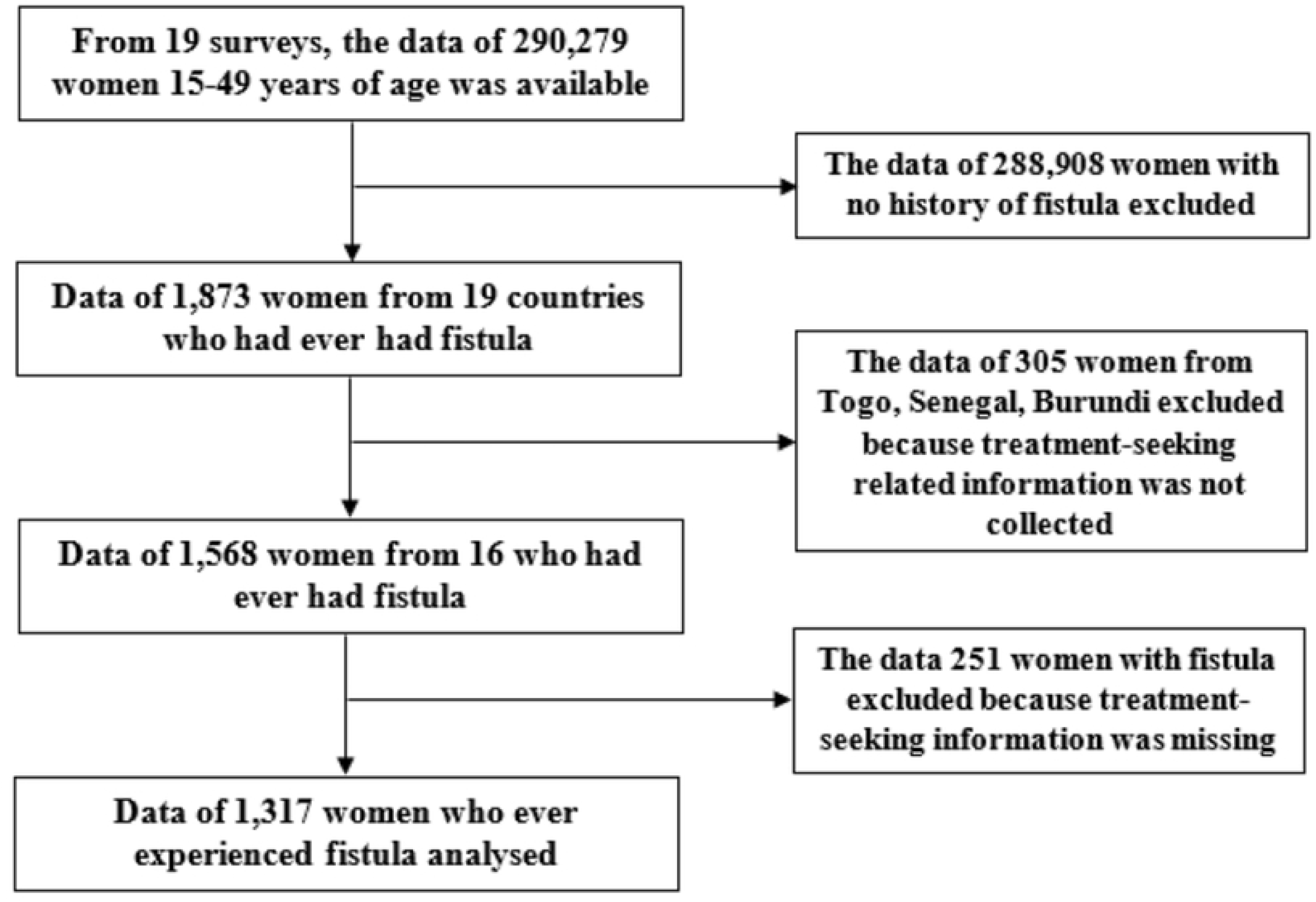
Flow chart of the study

### Sample size and sampling technique

As the study was conducted based on secondary data, priory sample size determination was not made. However, the available number of cases (n=1,317) is considered sufficient to determine the proportion of treatment seeking for vaginal fistula (68%) with 95% confidence level, 4% margin of error and design effect of 2. Country-wise, the unweighted sample size varied from 13 in Burkina Faso to 246 in Uganda (Supplementary file 1).

The DHS identified eligible subjects (women 15-49 years of age) using two-stage multiple cluster sampling approach. Initially a random sample of primary sampling units (enumeration areas) were drawn and in each unit 20-30 households were randomly selected. Ultimately, all eligible women in the selected households, irrespective of their marital status or birth experience, got interviewed [13].

### Data collection

The primary data were collected by trained enumerators, supervisors and data editors using pretested and standardized questionnaires prepared in the major local languages of the respective countries [13]. In all the surveys the presence of vaginal fistula was assessed by asking all women the question “did you experience a constant leakage of urine or stool from vagina?”. Then self-reported cause of the leakage was explored and classified as: pregnancy-related, sexual assault, pelvic surgery complications and others. Treatment-seeking practice and reasons for not seeking care were explored. In most of the surveys, follow up questions regarding whether the problem followed normal or difficult prolonged labour, whether the delivery ended up in live or stillbirth, were explored. Fistula-related survey questions forwarded in the 16 surveys are summarized provided as a supplementary file (Supplementary file 2).

### Data management and analysis

The datasets of the 16 surveys were downloaded from the measure DHS website and merged into one. Information about women who never experienced fistula was excluded and the remaining data were further cleaned and recoded as needed. Data were analysed using the survey data analysis approach via STATA software. Sampling weights provided in the original datasets and post-stratifications weights developed based on the 2017 population size of the countries were used for weighting. Wealth index was calculated as the indicator of economic status based on ownership of selected household assesses using Principal Component Analysis. The association between treatment-seeking and basic socio-demographic characteristics (age, place of residence, educational status, wealth index and marital) was assessed using mixed-effects logistic regression model and the outputs are presented using crude (COR) adjusted odds ratio (AOR) with the respective 95% confidence intervals (CI). The analysed dataset is provided as a supporting file (Supplementary file 3)

### Ethical considerations

Ethical clearance has not been sought for this specific analysis. However, the original surveys had been approved by the Institutional Review Board of ORC Macro and national-level ethical committees of all countries. We accessed the datasets after securing permission from the Measure DHS.

## Results

### Socio-demographic characteristics

The data of 1,317 women who ever experienced virginal fistula were analysed. While 16 countries had been represented in the dataset, more than half (60.0%) of the women were from six countries: Uganda, Chad, Malawi, Benin and Sierra Leone. The mean (± SD) age of the women was 30.7 (± 8.9) years and two-thirds (66.0%) were younger than 35 years. The majority were rural inhabitants and had no or limited formal education. In terms of marital status, 13.7% were widowed, divorced or separated. Intense reproductive experience appears to be common among the women as nearly one-third (31.2%) had already given birth five or more children. Further, 40.8% of women were first married before 18 years of age (Table 1).

**Table 1:** Socio-demographic characteristics of women who ever experienced vaginal fistula, 16 sub-Saharan Africa countries, 2010-2017.

| Characteristics (n=1317) | Frequency | Percentage |
| --- | --- | --- |
| Place of residence |  |  |
| Urban | 493 | 37.4 |
| Rural | 824 | 62.6 |
| Age in years |  |  |
| 15-24 | 367 | 27.9 |
| 25-34 | 502 | 38.1 |
| 35-49 | 448 | 34.0 |
| Educational status |  |  |
| No formal education | 456 | 34.6 |
| Primary | 535 | 40.6 |
| Secondary | 276 | 20.9 |
| Higher | 50 | 3.8 |
| Marital status |  |  |
| Never in union | 165 | 12.5 |
| Married/living with partner | 972 | 73.8 |
| Widowed | 65 | 5.0 |
| Divorced | 43 | 3.2 |
| Separated | 72 | 5.5 |
| Total children ever born |  |  |
| 0 | 153 | 11.6 |
| 1-2 | 428 | 32.5 |
| 3-4 | 325 | 24.7 |
| 5 or more | 411 | 31.2 |
| Age at first marriage (n=1211) |  |  |
| Before 18 years | 475 | 40.8 |
| 18 years or older | 689 | 59.2 |

**Table 2:** Reported causes of vaginal fistula, 16 sub-Saharan Africa countries, 2010-2017.

| Causes of fistula | Frequency | Percent |
| --- | --- | --- |
| Followed delivery | 887 | 67.6 |
| Sexual assault | 50 | 3.8 |
| Pelvic surgery | 36 | 2.7 |
| Unspecified non-obstetric causes | 87 | 6.6 |
| Others (unspecified) | 64 | 4.9 |
| Don't know | 188 | 14.3 |

### Causes of fistula

Nearly two-thirds (67.6%) of the women reported that the problem of leakage of urine or stool from vagina followed delivery/childbirth implying obstetric fistula. Smaller proportions identified sexual assault (3.8%) and pelvic surgery (2.7%) as the causes of the leakage. In the remaining 25.8% of the cases responses like “don’t know”, “don’t know but did not followed pregnancy” and “others” were given, hence clear-cut causes could not be ascertained. Excluding these ambiguous causes, 91.2% (887 out of 972) of the cases had obstetric fistula.

Most of the surveys included in the analysis presented additional information on obstetric fistula including time of onset of the leakage and whether the problem followed normal or difficult prolonged labour. Accordingly, about a quarter (23.3%) of the leakage started immediately or within the first day of birth while nearly half (46.6%) developed it late after one week of birth. In 57.4% of the cases the leakage followed prolonged or very difficult labour and 14.1% of the index pregnancies ended up in stillbirths (Table 3).

**Table 3:** Characteristics of obstetric fistula, 16 sub-Saharan Africa countries, 2010-2017.

| Characteristics of obstetric fistula | Frequency | Percent |
| --- | --- | --- |
| Onset of leakage (n=529) |  |  |
| Immediately or within the first day of birth | 123 | 23.3 |
| 2-6 days after birth | 159 | 30.1 |
| 1-6 weeks after birth | 157 | 29.6 |
| After the first 6 weeks of birth | 90 | 17.0 |
| Was the baby born alive (n=788) |  |  |
| Yes | 663 | 74.8 |
| No | 125 | 14.1 |
| Problem started after normal or difficult labour? (n=538) |  |  |
| Normal labour | 229 | 42.6 |
| Prolonged and very difficult labour | 309 | 57.4 |

### Health seeking for obstetric fistula

Of the women who had fistula, only 60.3% (95% CI: 56.9-63.6%) sought care from the formal medical sector and 57.6% received any modern treatment. Based on the available information in the dataset, in 1158 of the women it was possible to ascertain whether they had undergone fistula repair-surgery or not. It was found that only 28.5% (95% CI: 25.3-31.6%) women underwent surgery. A variety of reasons were reported for not seeking modern care. The leading were: lack of the awareness that fistula can be fixed (21.4%), don’t know where to go (17.4%), fear of cost associated with the treatment (11.9%), the problem resolved by itself (11.9%), embarrassment (7.9%) and inaccessibility of treatment centres (6.5%) (Table 4).

**Table 4:** Health seeking for vaginal fistula, 16 sub-Saharan Africa countries, 2010-2017.

| Variables | Frequency | Percentage |
| --- | --- | --- |
| Sought care from the formal medical sector (n=1317) |  |  |
| Yes | 793 | 60.3 |
| No | 523 | 39.7 |
| Person last sought treatment from (n=793) |  |  |
| Health professionals at health facilities | 767 | 96.6 |
| Community health workers | 27 | 3.4 |
| Received any modern treatment for the fistula (n=1317) |  |  |
| Yes | 759 | 57.6 |
| No | 558 | 63.4 |
| Underwent surgery for the fistula (n=1158) |  |  |
| Yes | 329 | 28.5 |
| No | 829 | 71.5 |
| Reasons for not seeking care (n=468) |  |  |
| Doesn't know fistula can be fixed | 100 | 21.4 |
| Do not know where to go | 81 | 17.4 |
| Treatment too expensive | 84 | 17.9 |
| Problem disappeared by itself | 56 | 11.9 |
| Embarrassment | 37 | 7.9 |
| Treatment facility too far | 31 | 6.5 |
| Could not get permission | 10 | 2.1 |
| Poor quality of care | 6 | 1.2 |
| Unspecified other reasons | 33 | 7.1 |

### Effectiveness of fistula-repair surgery

Among 244 women who underweight fistula-repair surgery, the effectiveness of the treatment was assessed. Three-quarters (74.1%) reported the leak completely stopped after surgery while the remaining reported that it did not stop but reduced (24.3%) or not stopped at all (1.6%). In aggregate, complete continence after surgery was not achieved in 25.9% of the cases.

### Health seeking behaviour and socio-demographic characteristics

Table 5 presents pattern of health seeking behaviour disaggregated by basic socio-demographic characteristics including current age, place of residence, educational status household wealth index and marital status. Among teenage girls with fistula only 31.5% sought care and as compared to adults 35 years or above, the odds of seeking care were reduced by 69% [AOR=0.31 (95 % CI: 0.20-47)]. Similarly, as compared to women who had any level of formal education, the odds of seeking care were reduced by 31% among women devoid of formal education. No meaningful differences were observed across categories of household wealth index, place of residence and current marital status (Table 5).

**Table 5:** Health seeking behaviour disaggregated by basic socio-demographic variables, 16 sub-Saharan Africa countries, 2010-2017.

| Variables | % who sought care | Odds Ratio |  |
| --- | --- | --- | --- |
|  |  | Crude | Adjusted |
| Current age (years) |  |  |  |
| 15-19 | 31.5 | 0.36 (0.24-0.54)* | 0.31 (0.20-0.47)* |
| 20-34 | 63.4 | 0.90 (0.71-1.16) | 0.86 (0.67-1.11) |
| 35 or above | 63.2 | 1 | 1 |
| Household wealth index |  |  |  |
| Poorest or poorer | 56.9 | 0.75 (0.58-0.97)* | 0.90 (0.66-1.21) |
| Middle | 60.3 | 0.79 (0.58-1.08) | 0.85 (0.60-1.20) |
| Richer or Richest | 61.4 | 1 | 1 |
| Current educational status |  |  |  |
| No formal education | 51.8 | 0.75 (0.57-0.98)* | 0.69 (0.51-0.93)* |
| Any formal education | 64.7 | 1 | 1 |
| Current place of residence |  |  |  |
| Urban | 62.6 | 1 | 1 |
| Rural | 58.9 | 0.73 (0.57-0.94)* | 0.82 (0.60-1.11) |
| Marital status |  |  |  |
| Married/living together | 61.3 | 1 | 1 |
| Others | 57.2 | 0.90 (0.70-1.16) | 1.00 (0.75-1.32) |

## Discussion

This secondary data analysis indicated that less than two-thirds of all vaginal fistula cases in SSA sought modern treatment and about a quarter underwent repair surgery. Treatment-seeking is constrained by multiple reasons including lack of awareness about the existence of treatment, embarrassment, financial and geographical inaccessibility to treatment centres. Further, health seeking is exceptionally low among teenagers and woman devoid of formal education. The study also found that in a quarter of women who already had fistula-repair surgery complete continence has not been attained.

We found that only 60% of fistula patients sought treatment from the formal medical sector and smaller proportions (29%) underweight surgery. The gap between the two figures suggests that the existing medical system is not capable of providing repair surgeries even to known cases. Previous studies in the region reported that the system fails to provide prompt treatment to registered cases due to lack of treatment centres, scarcity of skilled professionals and existence huge backlogs awaiting treatment [14-16]. A study in Somalia indicated 45% of women could not be able to undergo surgery within one year of registration [14] and in Burkina Faso women had to wait as long as five years to get operated [16].

We found that the leading reasons for failing to seeking treatment for fistula include lack of the awareness about the existence of treatment or where to get the treatment, economic constraints, embarrassment and inaccessibility of treatment centres. Likewise, a systematic review identified psychosocial, cultural, awareness, financial, and transportation domains as the critical barriers to fistula treatment in low income countries [12]. A study based on data from 20 countries concluded physical and financial inaccessibility of treatment cites and low awareness about existing treatment options as the key barriers to treatment seeking [16].

Unequivocal evidence exists that teenage pregnancy is associated with increased risk of obstetric fistula [17]. This study has indicated that treatment seeking is also exceptionally low (32%) among teenagers living with fistula. To the best of our knowledge, no study has explored the relationship between age and treatment-seeking for fistula. However, multiple studies have witnessed that adolescents are less likely to seek for basic maternity services than their adult counterparts due to multiple factors including lower socio-economic status, limited media exposure, influence from others and rural place of residence [18,19].

According to the WHO’s proposed indicators for monitoring and evaluating the quality fistula-repair surgery, closure rate should be 85% and complete continence should be achieved in 90% of women with a closed fistula [9]. Yet, we found that only 74% of women attained complete continence after surgery. The lower success rate may suggest the sub-optimal quality of the surgical care or presence of other contextual factors that limit surgical closure (e.g malnutrition) in the region. Facility-based studies conducted in Ethiopia, Rwanda, Nigeria and Guinea reported complete continence rates between 83 and 89% [20-23]. However, the figures cannot be directly compared to each other due to multiple reasons including differences in treatment success ascertaining approaches. Further, most facility-based studies evaluate closure rate at discharge but the current study accounted for treatment failure or recurrence that occurred any time after discharge.

One key limitation of the study is that presence of vaginal fistula and treatment outcome are determined based on self-reports of the women without any further clinical assessment. Therefore, misclassification bias cannot be excluded and have resulted in over-or underestimation the figures. However, a recent study in Nigeria has suggested, as compared to medical screening for fistula, self-reported symptoms have reasonably high sensitivity (92%) and specificity (83%) for identifying women affected by fistula [24].

Another limitation is that the DHS questionnaire measures lifetime occurrence of fistula and does not specifically tell when the incident actually happened. Consequently, indicators like treatment-seeking practice and surgical success rate may not exactly gauge the contemporary situations in the region. Accordingly, assuming access to treatment and quality of surgical care are improving, the study might have underrated the aforementioned two indices. Furthermore, measurement of the association between current socio-demographic variables (e.g. current educational status and wealth status) and treatment-seeking which happened sometime in the past bears an inherent erroneous assumption that socio-demographic variables are fixed and this leads to an obvious underestimation of the strength of association.

Finally, caution should be taken while generalizing the findings of the study to the entire SSA region because the health system and socio-demographic profile of the represented 16 countries may not be exactly the same as that of the remaining countries in the region. Additionally, many DHS conducted between 2010 and 17 in the region did not collected fistula-related information partly because the countries have not considered fistula as their priority problem. This is an additional evidence that the represented 16 countries are systematically different from the remaining SSA countries. However, we believe the study provides a reasonable picture of the treatment-seeking behaviour in the sub-continent.

## Conclusion

In the SSA region, treatment-seeking for fistula is low and is constrained by multiple factors – both from the health system and the patients’ perspectives. Women living with fistula have low treatment-seeking behaviour due to lack of awareness, sense of embarrassment as well as financial and physical inaccessibility to treatment centres. Further, the existing medical system is incompetent even to provide repair surgery to known cases. Teenage girls and women devoid of formal education are less likely to seek care. In the region, treatment-seeking should be augmented via a mix of strategies for abridging health-system, psycho-social, economic and awareness barriers.

## Acknowledgement

The authors acknowledge Measure DHS for granting access to the datasets.

## Supporting information captions

Supporting file 1: Sample size distribution by country.

Supporting file 2: Fistula-related survey questions forwarded in the 16 Demographic and Health Surveys.

Supporting file 3: Dataset analysed for the study.

